# Efficient mRNA Delivery *In Vitro* and *In Vivo* Using a Polycharged Biodegradable Nanomaterial

**DOI:** 10.1101/2024.11.05.622171

**Authors:** Xuejin Yang, Jingya Xiao, Daryl Staveness, Xiaoyu Zang

## Abstract

As RNA rises as one of the most significant modalities for clinical applications and life science research, efficient tools for delivering and integrating RNA molecules into biological systems become essential. We hereby report a formulation using a polycharged biodegradable nano-carrier, N1-501, which demonstrates superior efficiency and versatility in mRNA encapsulation and delivery in both cell and animal models. N1-501 is a polymeric material designed to function through a one-step facile formulation process suitable for various research settings. Its capability for mRNA transfection is investigated across a wide range of mRNA doses and in different biological models, including 18 tested cell lines and mouse models. This study also comprehensively analyzes N1-501’s application for mRNA transfection by examining factors such as buffer composition and pH, incubation condition, and media type. Additionally, N1-501’s superior *in vivo* mRNA transfection capability ensures its potential as an efficient and consistent tool for advancing mRNA-based therapies and genetic research.

## 1. Introduction

Gene therapy represents a revolutionary approach in medicine, offering immense potential for preventing and treating a wide array of diseases, including genetic disorders and cancer. The approval of several cell and gene therapy drugs worldwide has validated this potential, marking significant milestones in the field.[1,2] However, despite these advancements, the effective delivery of genetic cargo, particularly mRNA, remains a formidable challenge. mRNA cannot passively diffuse across cell membranes and is highly susceptible to degradation by extracellular nucleases without adequate protection.[3] Consequently, the development of efficient and safe gene delivery systems has become a critical focus in the field, attracting extensive research efforts.[4,5] Overcoming these delivery hurdles is imperative to fully harness the therapeutic potential of gene therapy and to expand its applications in clinical settings.

Viral delivery vehicles, such as adeno-associated viruses (AAV) and recombinant AAV (rAAV), have been widely utilized in gene therapy for both research and clinical applications.[6,7] However, the field faces notable challenges, such as immunogenic responses and the potential risk of insertional mutagenesis.[6,8] Additionally, the limited gene delivery capacity of viruses and the high cost of producing engineered viruses at large scale have further restricted their use as promising vectors.[9,10] Non-viral delivery systems have emerged as promising alternatives due to their lower immunogenicity and safer profiles. Current market transfection reagents, such as Lipofectamine 2000, Lipofectamine 3000, Lipofectamine CRISPRMAX, Lipofectamine RNAiMAX, FuGENE, and JetMESSENGER, have demonstrated advantages in *in vitro* applications. There have been continuous efforts to explore their application in delivering plasmid,[11,12] mRNA,[13] single-stranded oligonucleotides,[14] siRNA,[15] and gene-editing tools,[16] alongside efforts dedicated to optimizing formulation methodologies.[17–19] However, the performance of these commercially available reagents for *in vivo* use cases often falls short,[20,21] necessitating the development of more versatile and efficient transfection reagents that can function effectively in both *in vitro* and *in vivo* [22,23] applications. This is crucial to support the growing demands from both academic and industrial communities.

To fulfill the need for an efficient gene delivery method, a series of polymeric materials were developed, among which N1-501 was selected as an emerging tool in many original RNA research works (unpublished), representing an innovative advancement in non-viral gene delivery systems. N1-501 is designed to achieve high transfection efficiency with a straightforward formulation process, suitable for both *in vitro* and *in vivo* applications (Figure 3). In this study, the mRNA delivery efficacy of N1-501 across 18 cell lines is quantitatively assessed using eGFP expression assays, demonstrating versatile and robust performance across cell types. Importantly, systemic *in vivo* delivery of mRNA using N1-501 resulted in remarkable spleen tropism. Furthermore, the compatibility of different buffers, optimal incubation times, and precise formulation conditions for N1-501/mRNA nanoparticles were investigated. By fine-tuning parameters such as mRNA dose and transfection reagent dose, we aim to determine the operating window and optimize transfection efficiency and cell viability.

In conclusion, this study delves into the optimization strategies for mRNA transfection using N1-501, presenting comprehensive data on its performance across various conditions and applications. The dual utility of N1-501 for both *in vitro* and *in vivo* applications simplifies the translation from early-stage laboratory research to pre-clinical studies. This feature not only reduces costs but also accelerates the process of gene therapy development, addressing a critical challenges in the field. Overall, the study underscores the expansive potential of N1-501 and provides valuable insights into the enhancement of gene delivery systems, meeting the need for a versatile transfection reagent suitable for both laboratory and clinical settings.

## 2. Materials and Methods

### 2.1 Chemicals, Reagents, and Instruments

All chemicals and solvents were acquired from commercial sources (Sigma-Aldrich, VWR, Fisher, TCI, Oakwood Chemicals, ChemScene) and used without purification unless otherwise noted.

All consumables such as syringes, needles, syringe filters, glass vials, etc. were acquired from commercial sources (VWR, Fisher, Zoro, Chemglass).

All reagents used for the *in vitro* and *in vivo* assays were acquired from commercial sources as listed below: Dulbecco’s Modified Eagle Medium (DMEM) (C11995500CP, Gibco), Opti-MEM (51985-034, Gibco), 0.25% Trypsin-EDTA (25200-072, Gibco), fetal bovine serum (FBS) (S711-001S, Lonsera), PBS (21-040-CV, Corning), penicillin-streptomycin solution (C0222, Beyotime), Lipofectamine 3000 (L3000-001, Invitrogen), RNase-free water (R1600, Solarbio), CCK-8 (K1018, APExBIO), Hoechst 33342 (B2261, Sigma), eGFP mRNA (CT060, Catug Bio), recombinant human fibronectin (40113ES03, Yeasen), Fluc mRNA (Trilink L-7602), and IVISbrite D-Luciferin potassium salt (PerkinElmer cat. 122799).

Instruments used in this study included Varioskan® Flash (3001-1375, Thermo Scientific), High-Content Screening System (ImageXpress®Micro XL, Molelular Devices), Inverted Routine Microscope (TS100, Nikon), IVIS Lumina X5 imager (Perkin Elmer), and Living Image Software (PerkinElmer), Zetasizer Pro (Malvern).

### 2.2 In Vitro eGFP mRNA Transfection

#### 2.2.1 Cell Culture

The cell lines used in this study, including HEK 293T, HeLa, A549, MCF-7, HepG2, MIA PaCa-2, SH-SY5Y, DU 145, BT-474, Caco-2, NIH/3T3, MDA-MB-468, SW620, RAW264.7, NCI-H460, L929, U-87 MG, CHO-S were cultured following standard protocols. The cells were maintained at 37 °C in a 5% CO_2_ atmosphere. Detailed information on culture media, seeding density, and the commercial source of the cells can be found in Table S1.

#### 2.2.2 Cell Line Screening

Adherent or nonadherent cells were seeded approximately 24 hours before transfection using complete growth media. For example, HEK 293T cells were seeded at 15,000 cells/well in 100 µL DMEM supplemented with 10% (v/v) FBS and 1% (v/v) penicillin-streptomycin in black-walled 96-well plates pre-coated with recombinant human fibronectin. Cells were incubated 18–24 h at 37 °C (5% CO_2_) prior to transfection ensuring that confluency reaches 60%–80%. Right before transfection, the serum-containing media in each well were removed, and the cells were washed once with 100 μL of serum-free DMEM media per well. Then 50 μL of Opti-MEM was added to each well.

N1-501/mRNA complexes were prepared using a combination of 60 ng mRNA and 0.30 µL N1-501 stock solution (2.5 mg/mL in RNase-free water) per well. The complexes were formulated using a shaker mixing method. First, N1-501 solution was diluted in RNase-free water, followed by addition of mRNA stock solution (0.1 mg/mL in 1X PBS buffer) to achieve a final mRNA concentration of 0.01 mg/mL. The mixture was briefly pipetted 5 times at each step and further mixed on a shaker at 900 rpm for 30 seconds at 25 °C. After incubation at room temperature for 3 minutes, the sample was diluted with Opti-MEM [50 µL – 10 × V(mRNA)] and dispensed to each well, reaching a total volume of 100 μL per well for treatment. The Lipofectamine 3000 control was prepared in Opti-MEM according to the manufacturer’s instructions. Six replicates were run for all conditions. After 4-hour incubation at 37 °C, 100 µL of complete media was added to each well. The transfection efficiency and eGFP expression levels were evaluated using a high-content screening system 24 hours post-transfection. The data presented are the average percentage of GFP-positive cells and mean fluorescence intensities, with error expressed as ± SD. Other cell lines were treated in the same manner as HEK 293T cells.

#### 2.2.3 Design Space Exploration

The studies were conducted using HEK 293T cells. The sample preparation follows the same procedure as described in section 2.2.2, using various combinations of mRNA dose (15, 30, 60, 90, 120 ng per well) and N1-501 dose (0.06, 0.10, 0.12, 0.15, 0.18, 0.20, 0.24, 0.30, 0.35, 0.40, 0.50, 0.60, 0.70 µL per well). The Lipofectamine 3000 control was prepared in Opti-MEM according to the manufacturer’s instructions. All conditions were tested in at least triplicate, with duplicated groups independently formulated to ensure reproducibility and consistency. After 4-hour incubation at 37 °C, 100 µL of complete media was added to each well. The transfection efficiency and eGFP expression levels were evaluated using a high-content screening system 24 hours post-transfection. The data presented are the average percentage of GFP-positive cells and mean fluorescence intensities, with error expressed as ± SD.

#### 2.2.4 Buffer and Media Effect

The studies were conducted using HEK 293T cells. The sample preparation follows the same procedure as described in section 2.2.2, using a combination of 30 ng mRNA and 0.15 µL N1-501 stock solution per well. Samples were formulated in 10 mM Citrate buffer (pH = 4.0, 5.0, 6.0), 10 mM acetate buffer (pH = 4.0, 5.0, 6.0), 1X PBS buffer (pH = 7.4), 10 mM HEPES buffer (pH = 8.0), and water, respectively. These samples were further diluted in Opti-MEM and tested in HEK 293T cells in Opti-MEM. To explore media effect on mRNA transfection, samples formulated in water were diluted in serum-free DMEM [DMEM(-/-)], DMEM with 10% FBS [DMEM(10% FBS)], or DMEM containing 10% FBS and 1% penicillin/streptomycin [DMEM(+/+)], and then applied to cells in the same media. All conditions were tested in quadruplicate. After 4-hour incubation at 37 °C, 100 µL of complete media was added to each well. The transfection efficiency and eGFP expression levels were evaluated using a high-content screening system 24 hours post-transfection. The data presented are the average percentage of GFP-positive cells and mean fluorescence intensities, with error expressed as ± SD.

#### 2.2.5 Incubation Condition

The sample preparation follows the same procedure as described in section 2.2.2, using a combination of 30 ng mRNA and 0.15 µL N1-501 stock solution per well. The samples were incubated for different durations (5, 10, 15, 20, 30, 40, 60 minutes) at different temperatures (0, 25, 37 °C) before applying to cells. All conditions were tested in triplicate, with duplicated groups independently formulated to ensure reproducibility and consistency. After 4-hour incubation at 37 °C, 100 µL of complete media was added to each well. The transfection efficiency and eGFP expression levels were evaluated using a high-content screening system 24 hours post-transfection. The data presented are the average percentage of GFP-positive cells and mean fluorescence intensities, with error expressed as ± SD.

#### 2.2.6 N1-501’s Shelf Life

The sample preparation follows the same procedure as described in section 2.2.2, using a combination of 60 ng mRNA and 0.30 µL N1-501 stock solution per well. N1-501 stock solutions used in this study were stored at -20 °C, 4 °C, and 25 °C for up to 6 months. All conditions were tested in quadruplicate, with duplicated groups independently formulated to ensure reproducibility and consistency. After 4-hour incubation at 37 °C, 100 µL of complete media was added to each well. The transfection efficiency and eGFP expression levels were evaluated using a high-content screening system 24 hours post-transfection. The data presented are the average percentage of GFP-positive cells and mean fluorescence intensities, with error expressed as ± SD.

### 2.3 Cell Viability Assay

The impact on cell viability of N1-501/mRNA complexes was assessed using a CCK-8 viability assay. After conducting image analysis using the high-content screening system 24 hours post-transfection, 10– 15 µL of CCK-8 reagent was added to each well. Following a 2-hour incubation, the absorbance at 450 nm was measured using a plate reader. Cell viability was calculated by dividing the absorbance of treated cells by that of untreated cells.

*<h2>2.4 Dynamic light scattering (DLS) for size and zeta potential measurement*

The sample preparation follows the same procedure as described in section 2.2.2, using 300 ng mRNA. After a 3-minute incubation at room temperature, the sample was diluted with 1200 μL of water and mixed thoroughly for characterization using the Malvern Zetasizer Pro. A 150 μL aliquot was transferred to a clear cuvette to measure nanoparticle size and polydispersity, while 900 μL was transferred to a capillary cell for zeta potential measurement. All reported values represent the average of at least three trial runs, with error expressed as ± SD.

### 2.5 In Vivo FLuc mRNA Transfection

#### 2.5.1 Animal

All animal experiments were consistent with local, state, and federal regulations as applicable. Food and water were supplied *ad libitum*. Female BALB/c mice (Charles River, UK), aged 6−8 weeks, were randomly assigned to groups (n = 3) and housed in a fully acclimatized room.

#### 2.5.2 In Vivo Formulation

N1-501/Fluc mRNA (Trilink L-7602) complexes were formulated using a pipette mixing method. To a 1.5 mL microcentrifuge tube was added 579 µL RNase-free water and 20 µL Fluc mRNA (1 mg/mL solution in 1X PBS) at room temperature. The solution was mixed by pipetting 10 times, followed by the addition of 121 µL N1-501 (5 mg/mL stock solution in RNase-free water). The sample was further mixed by pipetting 30 times (approximately once per second). Then 80 µL 10X PBS was added, giving a total volume of 800 µL with a 1X PBS concentration. The solution was thoroughly mixed, and 200 µL of the mixture, containing 5 µg of Fluc mRNA and 30.25 µL of N1-501 stock solution per mouse, was injected intravenously into each of three female BALB/c mice through tail vein. The negative control group, containing only 5 µg of Fluc mRNA per mouse, was prepared following the same method.

#### 2.5.3 BLI Imaging

The biodistribution of N1-501/mRNA complexes in Balb/c mice was determined by imaging the above animals with luciferin at 3-, 6- and 12-hours post-injection using the IVIS Lumina X5 imager (PerkinElmer). A fresh stock solution of luciferin was prepared at 30 mg/mL in 1X PBS, and each mouse was injected intraperitoneally with 100 µL of the luciferin solution 10 minutes before imaging. Mice were anesthetized with isoflurane and imaged using Living Image software (PerkinElmer) with automatic imaging settings (exposure time of 20 seconds to 4 minutes, depending on signal strength). Whole body images were captured in both ventral and lateral positions at each time point. Then two mice at 6 hours and one at 12 hours were immediately sacrificed. Major organs, including the brain, heart, lungs, liver, spleen, stomach, intestines, kidneys, and skeletal muscle, were promptly excised and imaged *ex vivo*.

#### 2.5.4 Analytic Method

Living Image software was utilized to analyze the numerical pixel values and image data from the acquisition files for all experimental groups. The saved Living Image files were loaded for image display and analysis. Circular Regions of Interest (ROIs) were placed around each luminescent source, and the system automatically displayed data for all the ROIs created within the images.

### 2.6 Statistical Analysis

All statistical analyses were conducted using GraphPad Prism v10.1.0 (GraphPad Software) and JMP Pro 17 (JMP Statistical Discovery LLC). Statistical analysis between the two groups was performed using a two-tailed, unpaired Student’s *t* test. For comparing more than two groups, statistical analysis of one-way or two-way ANOVA followed by Tukey, Bonferroni or Dunnett’s multiple comparison test was applied. A *p*-value of less than 0.05 was considered statistically significant.

## 3. Results and Discussion

### 3.1 N1-501 shows superior efficiency for the delivery and transfection of mRNA in vitro than other commonly used transfection reagent

N1-501 is compatible with a diverse range of cell types. To evaluate its mRNA delivery performance, 18 commonly used research cell lines were selected, with Lipofectamine 3000 used as a comparison (Figure 1A, 1B). N1-501 exhibited excellent transfection efficiency in the epithelial and fibroblast cell lines tested (Table 1). Moderate transfection efficiency was observed in brain cell lines. Over 80% GFP-positive rate was observed with N1-501 in RAW264.7, a notably high delivery rate for an immune cell line, which tend to be recalcitrant to mRNA transfection. Remarkably, N1-501 demonstrated higher eGFP expression levels (Figure 1B and Figure S1) and faster expression rates (Figure S2) in all cell lines screened compared to Lipofectamine 3000. The impact of mRNA transfection on cell viability was evaluated after 24-hour treatments of N1-501/eGFP mRNA and Lipofectamine/eGFP mRNA complexes, compared to the mRNA-only group and untreated control cells. Cell viability with N1-501 material remained largely unaffected (>85%) among most carcinoma, brain, and fibroblast cell lines (Table S2). Lower cell viabilities, as observed with Caco-2, RAW264.7, and L929 cells, are probably associated with higher transfection efficiencies and superior GFP expression levels using N1-501 compared to Lipofectamine 3000.[14] However, the effect of transfection reagents on cell viability can vary depending on the specific cell type and its sensitivity. In our study, we did not observe a strong correlation between cell viability and transfection efficiency (Figure S3). Further optimization on mRNA and N1-501 doses is necessary to maximize efficacy while maintaining a safe cellular profile. Overall, N1-501 effectively transfected a variety of cell lines and outperformed Lipofectamine 3000 in most cases.

**Table 1.**
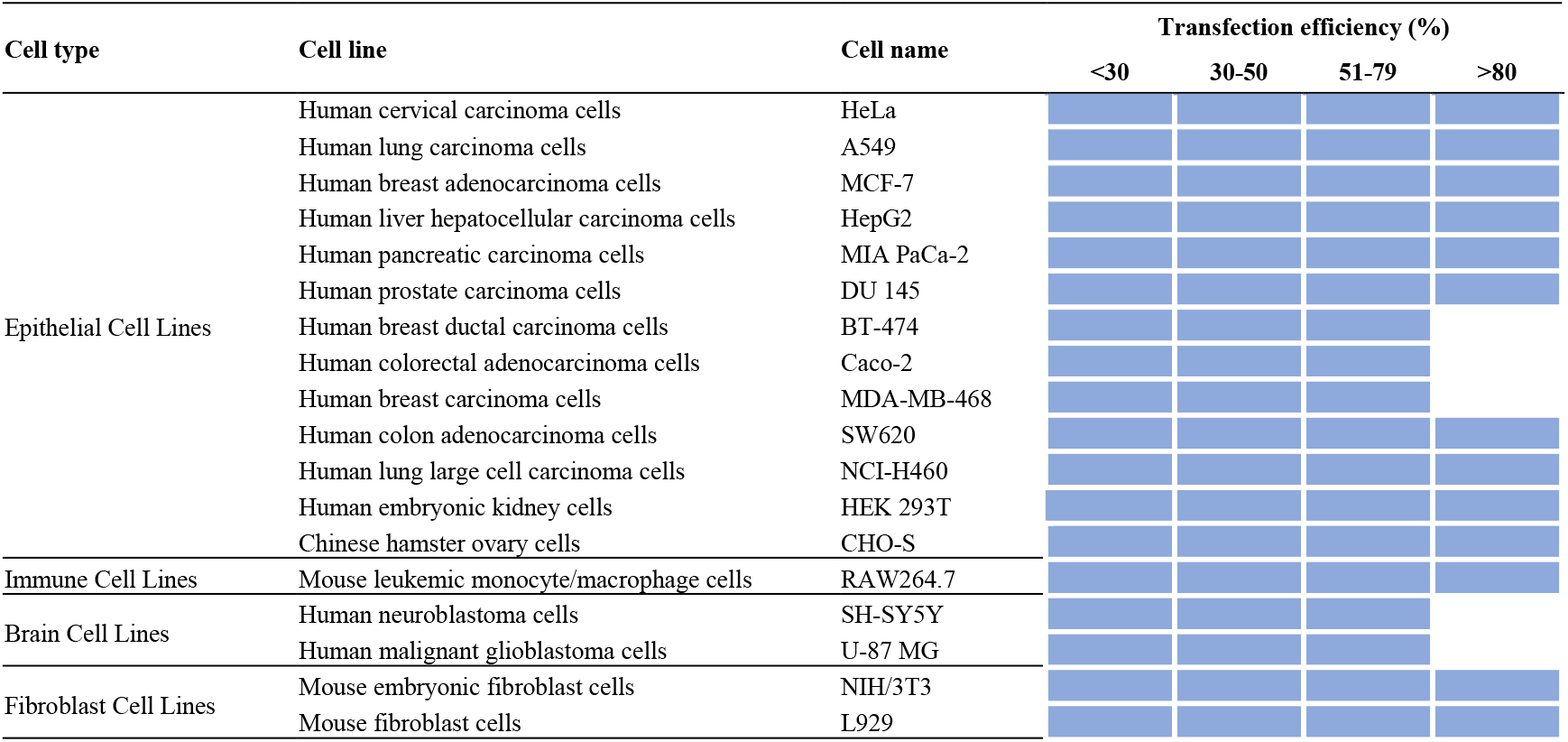
N1-501 delivers mRNA in a variety of cell lines.

**Figure 1.**
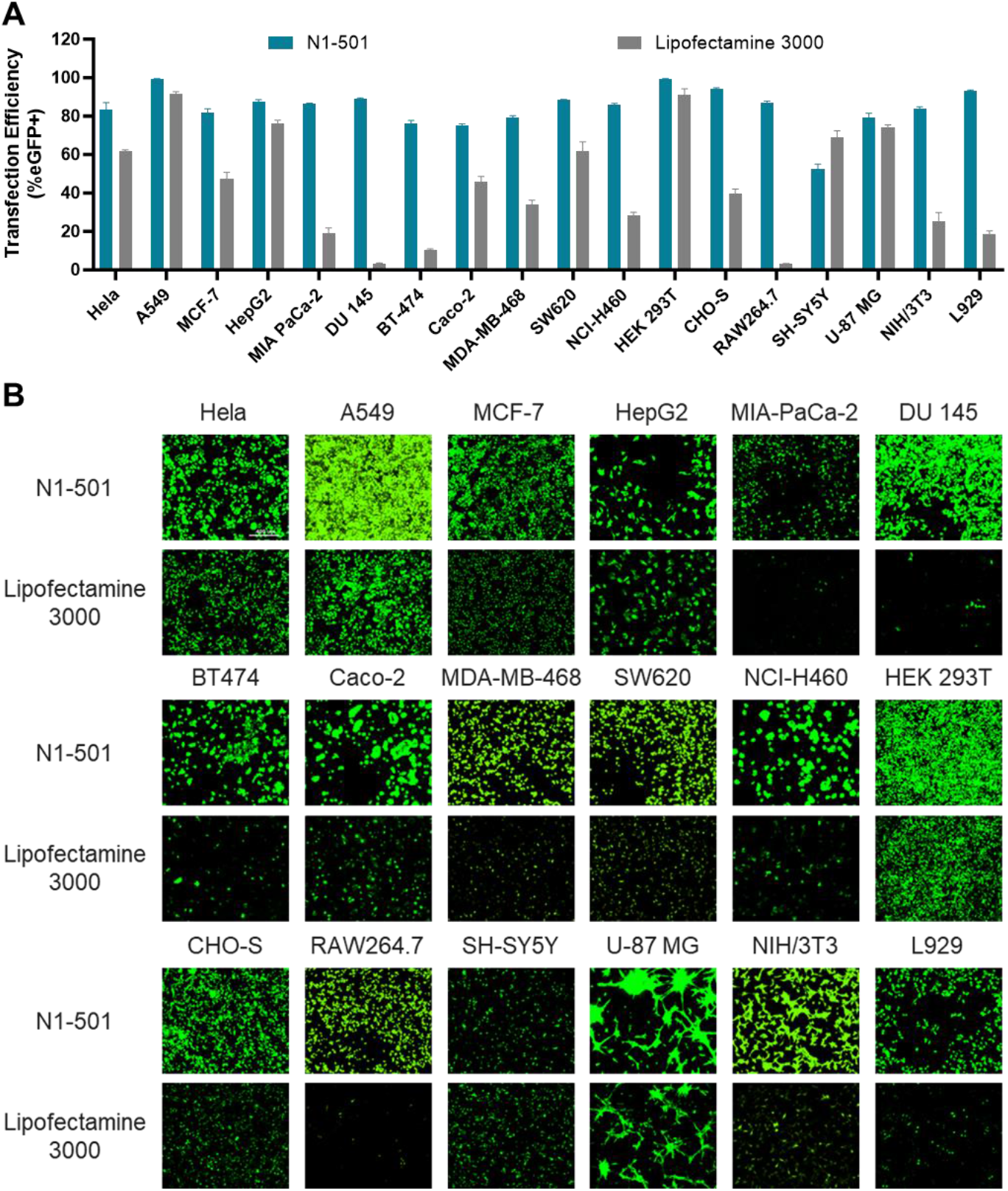
N1-501 effectively delivers mRNA across various cell lines as compared with Lipofectamine 3000. (A) The transfection efficiency was evaluated after 24 hours of treatment. (B) Fluorescence microscopy images of 18 cell lines were captured 24 hours post-transfection to visualize eGFP expression levels (Scale bar, 400 µm).

### 3.2 N1-501 shows efficient delivery and transfection of mRNA in vivo

N1-501 is a dual-function material that delivers mRNA both *in vitro* and *in vivo*. To assess its *in vivo* performance, N1-501 complexed with luciferase mRNA was injected intravenously (i.v.) to BALB/c mice. The formulation containing only luciferase mRNA was used as negative control. After 3, 6, and 12 hours of administration, D-luciferin was injected intraperitoneally, and luciferase protein expression was evaluated at each time point by whole-body imaging with an IVIS Lumina X5 imager (Perkin Elmer). In each group, two mice were sacrificed after 6 hours, and one mouse was sacrificed after 12 hours. All major organs, including brain, heart, lungs, liver, spleen, stomach, intestines, kidneys, and skeletal muscle, were immediately imaged *ex vivo*.

The treated group showed superior delivery selectivity towards spleen at all time points (Figure 2A), with the signal diminishing between 3 and 12 hours (Figure 2C). The *ex vivo* imaging further proved the organ and tissue selectivity of mRNA delivery. Luminescence signals were primarily detected in spleen, lungs, and liver, with the luminescence intensity in spleen being more than two orders of magnitude higher than that of the lungs and liver, elaborating N1-501’s spleen targeting capability for mRNA delivery (Figure 2B).

**Figure 2.**
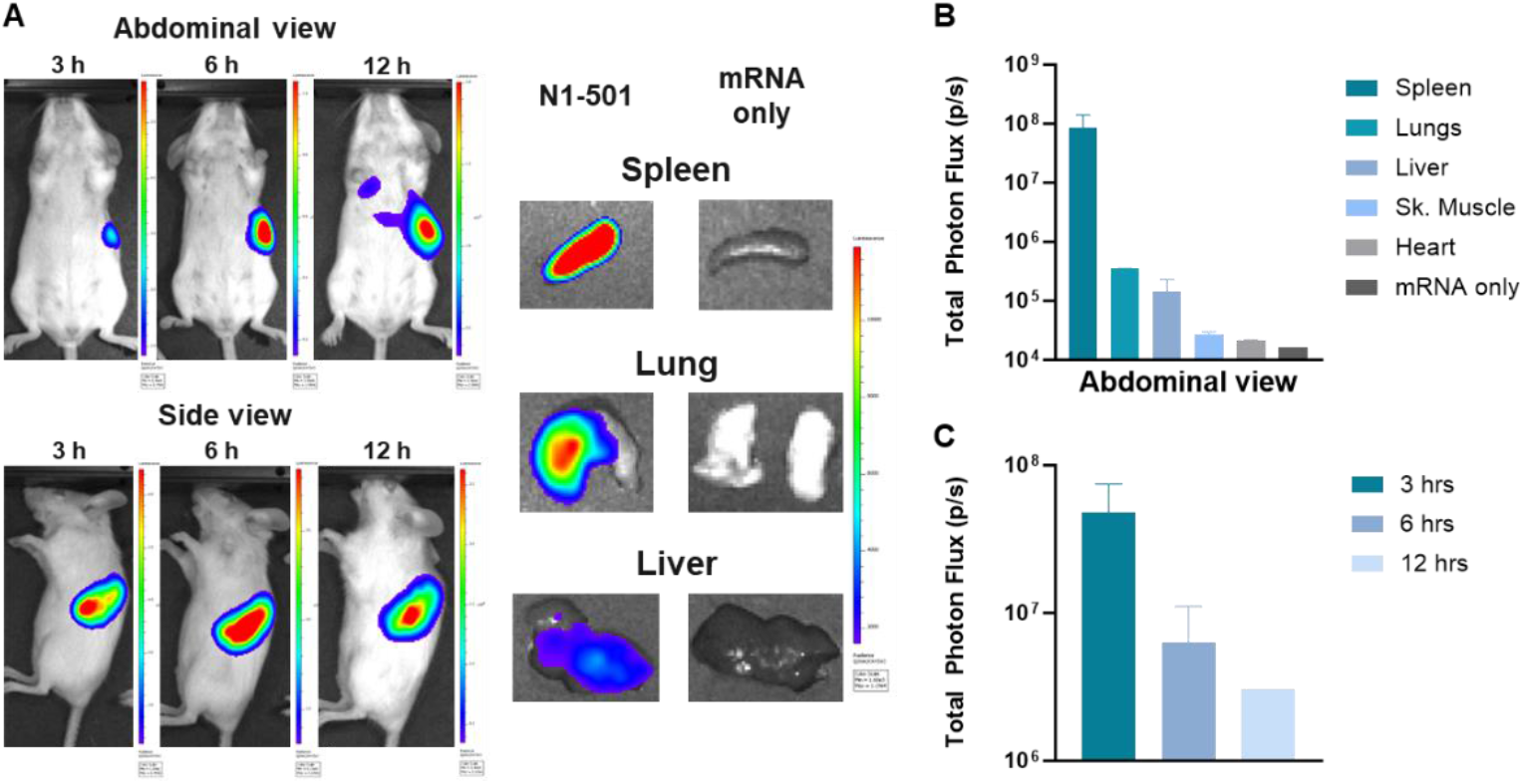
N1-501 demonstrated high spleen tropism during *in vivo* transfection. (A) *In vivo* bioluminescence (3, 6, 12 h) and *ex vivo* bioluminescence (6 h) of major organs excised from Balb/c mice IV injected with N1-501/Fluc mRNA. (B) Quantification of total bioluminescence in various organs. (C) Quantification of total bioluminescence in mice at different time points.

**Figure 3.**
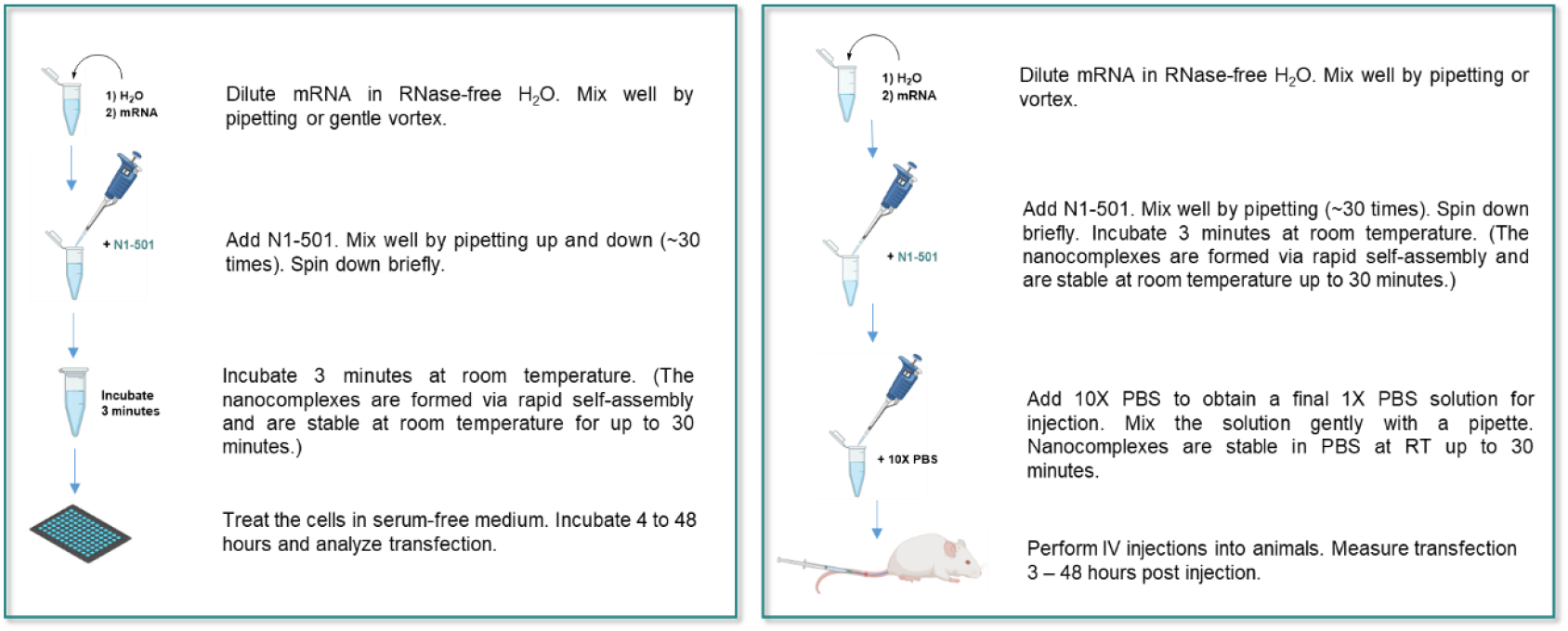
Graphical depiction of operationally simple transfection protocols developed for N1-501.

### 3.3 N1-501 functions through a one-step facile formulation process

N1-501 can be easily formulated with mRNA using a simple pipette mixing method, as illustrated in Figure Its water-soluble nature allows the entire formulation process to take place in an aqueous system, eliminating the use of any organic solvents or specialized buffer systems. Upon mixing N1-501 with diluted mRNA in RNase-free water, nanoparticles form through self-assembly, and the sample is ready for cell transfection after a 3-minute pre-incubation at room temperature. For *in vivo* applications, the sample should be adjusted to a 1X PBS solution before administering to animals.

Mixing the sample with a shaker offers better control over the formulation process on a small scale. In this approach, the N1-501 stock solution is first diluted in RNase-free water, followed by addition of mRNA stock solution. The sample is then mixed on a shaker at 900 rpm for 30 seconds under 25 °C. This shaker mixing method provides more consistent particle sizes compared to pipette mixing. Both methods are easily accessible and can be performed under standard laboratory conditions.

### 3.4 N1-501 offers a wide operational window for RNA formulation and delivery across an extensive dose range

Cell transfection rate, mRNA expression level, and cell viability are the three key criteria for evaluating mRNA transfection. For specific applications, RNA dosing can vary widely based on research objectives and experimental settings. To explore the safety window and functional range, the amount of mRNA dose and N1-501 dose were adjusted during the formulation process for initial screening. Figure 4 illustrates the design space of N1-501 at various combinations of mRNA and reagent doses using HEK 293T cell lines in a 96-well assay. A broad range of mRNA doses, from 15 ng to 120 ng per well, combined with N1-501 doses ranging from 0.06 to 0.7 µL per well, were tested, resulting in a matrix of 65 conditions. The GFP expression level, represented by mean fluorescence intensity (MFI), exhibited a dependence on mRNA dose. Increasing the mRNA dose, along with an increased N1-501/mRNA ratio, led to higher MFI (Figure 4A) but compromised the cell viability (Figure 4C). This suggests that in scenarios where a high mRNA dose is needed, a lower N1-501/mRNA ratio is necessary to balance efficacy and safety. Notably, the transfection efficiency remained above 96% across all mRNA doses (Figure 4B). The results can serve as a reference for researchers to identify boundary conditions and optimal ranges for their specific applications.

**Figure 4.**
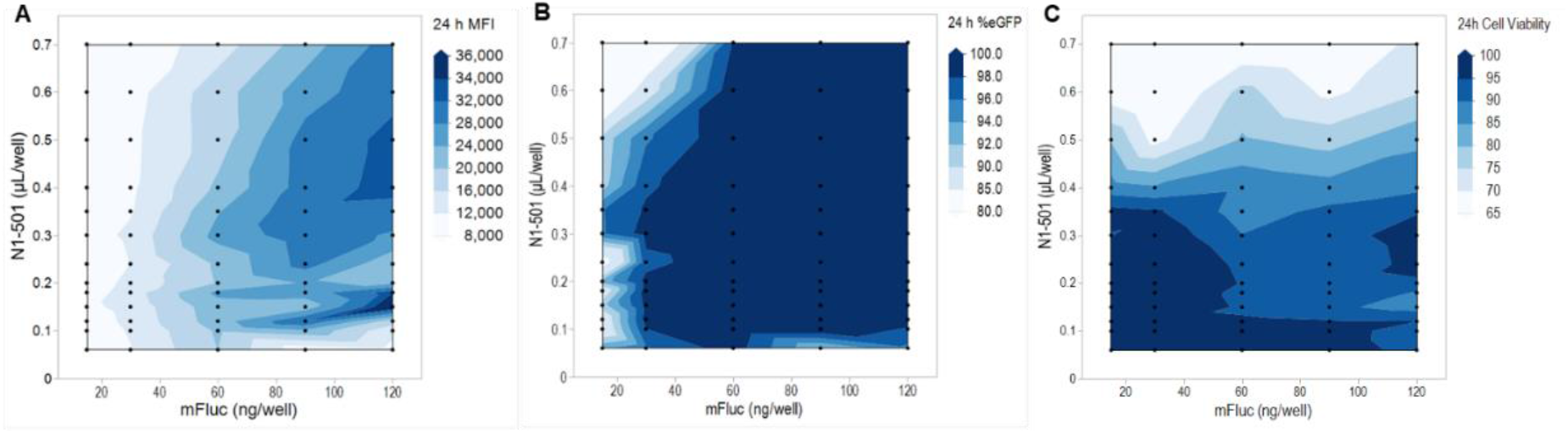
The transfection performance and safety profiles of N1-501 at varied mRNA and reagent dose combinations. Black dots represent specific formulation conditions used in the study; the contour plot was filled with simulated data. (A) The contour plot for GFP expression, represented by MFI, was measured using a high-content screening system 24 hours after transfection. (B) The contour plot showing the percentage of GFP-positive cells 24 hours post transfection. (C) The contour plot showing HEK 293T cell viability 24 hours after treatment with N1-501/eGFP mRNA complexes. Viability was evaluated using CCK-8 assay.

### 3.5 N1-501 is compatible with various buffer and medium conditions for different research purposes

Physiological buffers are widely used in cell culture, molecular biology experiments, animal studies, and clinical research. In gene delivery, they serve various roles, such as the storage of gene cargos, aiding in formulation process, serving as nanoparticle storage solutions, and acting as the formulation solution for *in vivo* administration.[24,25] Buffers with a specific pH range are crucial for maintaining the stability and functionality of gene cargos. They also affect the charge state of ionizable transporter materials, which in turn, influences nanoparticle encapsulation efficiency, size, uniformity, and stability. In this study, four common physiological buffers*—*citrate, acetate, phosphate-buffered saline (PBS), and HEPES*—*were tested during the formulation process, and their effects on transfection performance were evaluated *in vitro*.

Citrate and acetate buffers, each at a 10 mM concentration with sodium counterions, were adjusted to pH levels of 4.0, 5.0, and 6.0 to screen for acidic conditions. A 10 mM HEPES buffer at pH 8.0 was selected for basic condition, while 1X PBS buffer at pH 7.4 was chosen for neutral condition. N1-501 was diluted in each of these buffers or in RNase-free water and then mixed with mRNA. After a 3-minute pre-incubation, the nanoparticle solutions were diluted in reduced serum medium Opti-MEM and applied to cells in the same media. MFI (Figure 5A) and the percentage of GFP+ cells (Figure 5B) were measured 24 hours post-transfection. GFP expression levels showed a clear pH dependence, as nanoparticles formulated in citrate buffer at pH 6.0 produced the highest MFI, followed by pH 5.0 and pH 4.0 (Figure 5A). A similar trend was seen with acetate buffers. MFI in PBS at pH 7.4 and HEPES at pH 8.0 were comparable to that in water. Interestingly, the transfection efficiency showed no significant differences across all buffers and pH environments (Figure 5B). This indicates that the N1-501 is robust under various buffer conditions.

**Figure 5.**
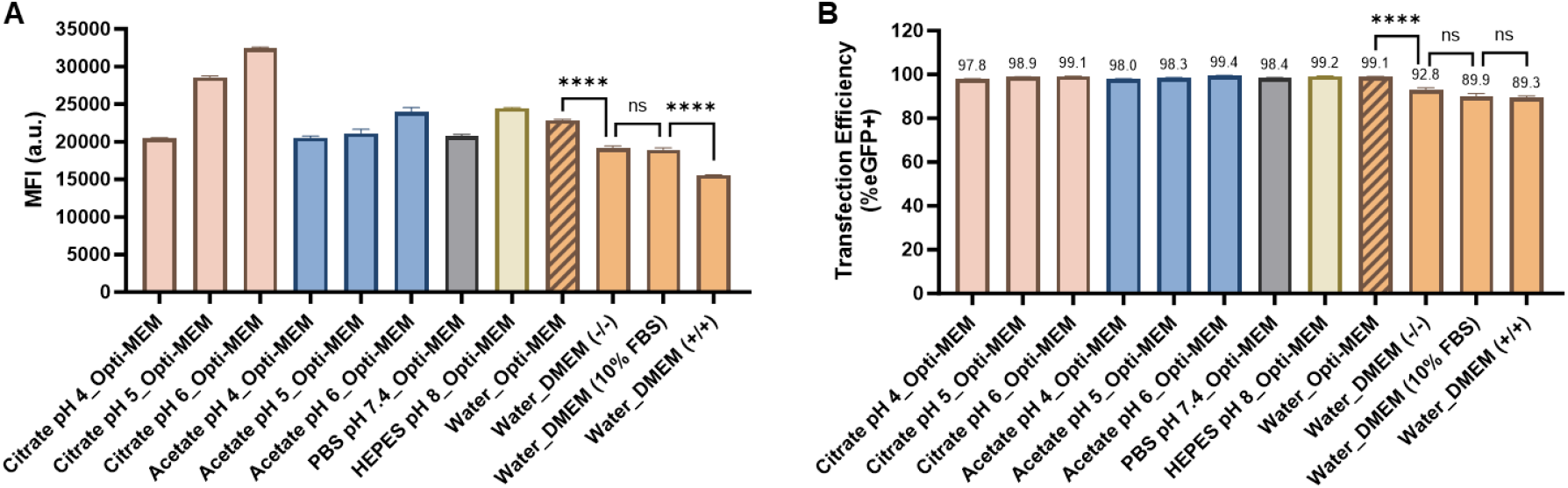
Buffer and media compatibility during formulation and transfection. N1-501 was diluted in 10 mM Citrate buffer (pH = 4.0, 5.0, 6.0), 10 mM acetate buffer (pH = 4.0, 5.0, 6.0), 1X PBS buffer (pH = 7.4), 10 mM HEPES buffer (pH = 8.0), and water, respectively. After mixing with mRNA, the nanoparticle was diluted in reduced serum medium Opti-MEM, serum-free DMEM [DMEM(-/-)], DMEM with 10% FBS [DMEM(10% FBS)], or DMEM containing 10% FBS and 1% penicillin/streptomycin [DMEM(+/+)] and treated to cells in the same media. The MFI (A) and percentage of GFP-positive cells (B) were assessed 24 hours after transfection. Data are shown as mean ± SD. Statistical significance was determined using an unpaired two-tailed Student’s *t* test (ns: no significance, \*\*\*\**p* < 0.0001).

The serum tolerance of N1-501 was also assessed. Serum can interfere with mRNA transfection by potentially introducing RNase contamination, competing with transfection vehicles for cell-surface receptors for entrance into cells[26], or destabilizing the mRNA-vehicle complex[27,28]. Therefore, serum-free or serum-reduced media (such as Opti-MEM) is often recommended for optimal transfection performance. However, it should be noted that this condition may not be suitable for applications and/or cell lines requiring continuous serum exposure. In this study, we tested whether N1-501 could effectively deliver mRNA in the presence of FBS. N1-501/eGFP mRNA, formulated in RNase-free water, was diluted with serum-free DMEM [DMEM(-/-)], DMEM with 10% FBS [DMEM(10% FBS)], or DMEM containing 10% FBS and 1% penicillin/streptomycin [DMEM(+/+)]. The samples were then treated to cells in the corresponding media. Compared to Opti-MEM, the transfection efficiency decreased in DMEM, while the effect of serum was not significant (Figure 5B). However, the media change and the presence of serum had a more pronounced impact on GFP expression levels, with a notable reduction in MFI observed in DMEM, DMEM(-/-), and DMEM(+/+) (Figure 5A). These findings suggest that N1-501 can successfully transfect cells in the presence of serum. The highest level of delivery occurs in serum-free or serum-reduced media in this experimental setup, but this data indicates that one has the flexibility to consider a range of media conditions while optimizing for a given application.

Cell viability of HEK cell line treated with N1-501/eGFP mRNA nanoparticles formulated in various buffers, pH ranges, and media was assessed using the CCK-8 assay after 24 hours of treatment. Except for DMEM(-/-) media, with a viability below 90%, all the other conditions exhibited over 94% cell viability, demonstrating the safety of N1-501 across a range of formulation and treatment conditions (Figure S4).

### 3.6 Formulation time and temperature do not affect the reproducibility of N1-501/mRNA transfection capability

The robustness and reproducibility of nanoparticle formulations are crucial for mRNA transfection experiments. Stable nanoparticles preserve mRNA integrity and simplify material handling and transportation when needed.[29–31] Additionally, it supports time-intensive tasks, such as high-throughput studies and situations where multiple experiments must be carried out concurrently. A stable and consistent formulation not only maintains transfection efficacy but also improves experimental success rates and ensures reliable, reproducible results.

In a typical *in vitro* or *in vivo* study, nanoparticles can either be prepared and used immediately or held until a matrix of samples is ready for treatment. In certain cases, samples may need to be transported to the animal facility for administration, requiring consideration of time and temperature during transport.

To assess the formulation robustness under standard laboratory conditions, the effect of incubation time (or pre-incubation time) on mRNA delivery by N1-501 was tested *in vitro*. Upon formulation, the samples were stored at 0 °C, 25 °C, and 37°C for different times before being applied to HEK 293T cells. After 24 hours of transfection, the performance of each condition was analyzed and compared. As shown in Figure 6B, no significant differences in transfection efficiency were observed within 60 minutes of formulation at 0 or 25 °C (>98%), and the GFP-positive rate remained above 95% after one hour of incubation at 37 °C. It should be noticed that the sample pre-incubated under different temperature and durations was formulated independently. The consistent high transfection efficiency across all conditions demonstrates the formulation’s excellent robustness and reproducibility, allowing it to accommodate the non-uniformity that can arise when performing high-throughput experiments, translational studies, or other complex research endeavors.

**Figure 6.**
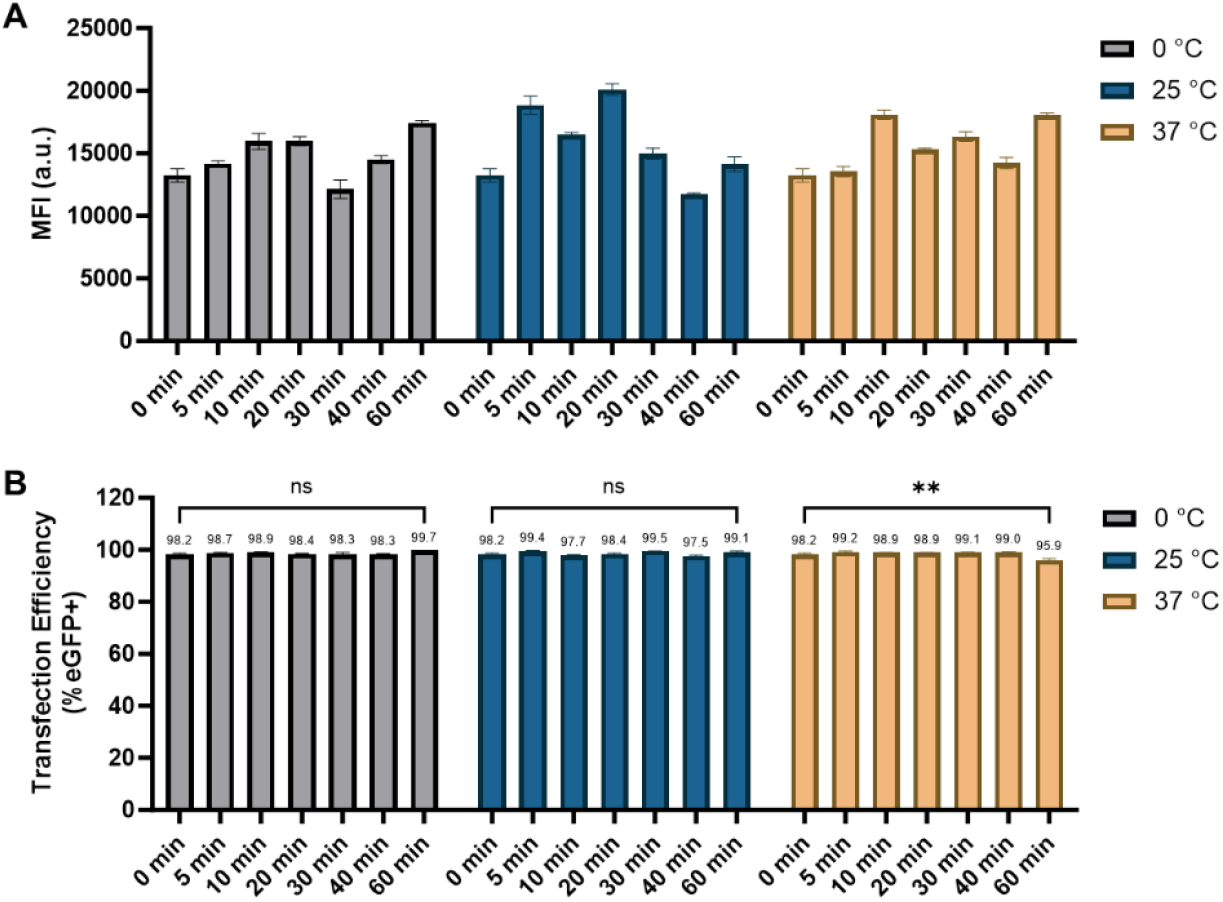
Effect of pre-incubation time on mRNA delivery by N1-501. N1-501 was formulated with eGFP mRNA and then held at 0 °C, 25 °C, and 37 °C for different times. (A) MFI and (B) percentage of GFP-positive cells were measured 24 hours after transfection. Data are shown as mean ± SD. Statistical significance was determined using an unpaired two-tailed Student’s *t* test (ns: no significance, \*\**p* < 0.01).

### 3.7 N1-501 is a shelf-stable tool for long-term research use

The shelf life of N1-501 was assessed based on its transfection efficacy *in vitro*. Its stock solution (2.5 mg/mL in RNase-free water), stored at -20 °C, 4 °C, and 25 °C for various time periods, was formulated with eGFP mRNA at a N1-501 volume (µL/well) to RNA weight (ng/well) ratio of 0.005 and tested in HEK 293T cells. On the day of transfection, an aliquot of N1-501 in solid state form was dissolved with RNase-free water (2.5 mg/mL) and used as fresh sample for comparison. As shown in Figure 7A and 7B, N1-501 stock solution retained its stability and potency for up to 6 months when stored at 4 °C or -20 °C. In contrast, storage at 25 °C resulted in a slight performance decrease after 2 months and a statistically significant reduction after 6 months (Figure 7A and Figure S5). Additionally, no noticeable influence on performance has been observed when stored as a solid at -20 °C for 8 months (data not shown).

**Figure 7.**
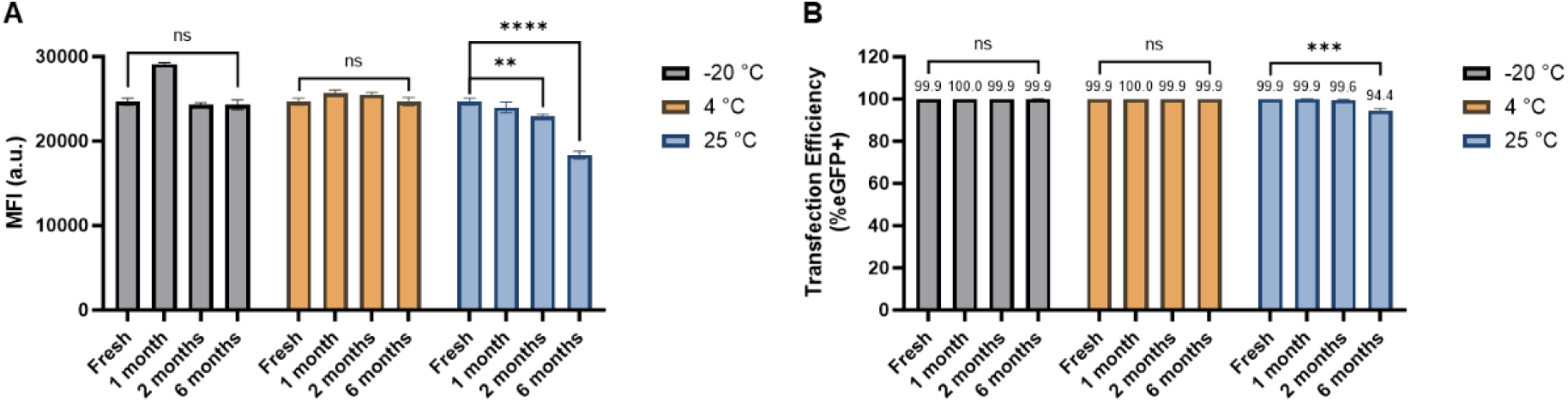
N1-501’s shelf life evaluated by transfection efficacy *in vitro*. (A) MFI and (B) percentage of GFP-positive cells were assessed 24 hours after transfection. Data are shown as mean ± SD. Statistical significance was determined using an unpaired two-tailed Student’s *t* test (ns: no significance, \*\**p* < 0.01, \*\*\**p* < 0.001, \*\*\*\**p* < 0.0001).

## 4. Conclusions

N1-501 has been shown functional characteristics valuable to a new RNA delivery tool, demonstrating versatility in both *in vitro* and *in vivo* applications. Its straightforward formulation process, easily executed under standard laboratory conditions, coupled with stability across various temperatures, ensures consistent and reproducible results while simplifying handling protocols. N1-501 demonstrates adaptability by being compatible with a wide range of physiological buffers across diverse pH levels and exhibiting serum tolerance during mRNA transfection. This feature amplifies its practical utility in complex biological environments. Notably, N1-501 achieves high transfection efficiency over an expansive mRNA dose range, enhancing its applicability across various delivery scenarios. Outperforming industry-standard Lipofectamine 3000 in most cases, it efficiently transfects a diverse array of cell lines, including epithelial, fibroblast, and brain cells, showcasing its broad-spectrum effectiveness. N1-501’s prowess extends to gene editing, demonstrating remarkable success in delivering Cas9 mRNA and sgRNA, achieving up to 96% editing efficiency *in vitro* (data not shown). Importantly, N1-501 exhibits superior spleen tropism following systemic mRNA delivery *in vivo*, a distinctive feature that sets it apart in the realm of transfection reagents capable of bridging laboratory and clinical applications. This unique combination of attributes positions N1-501 as a versatile and powerful tool, poised to accelerate research and development across the spectrum of RNA-based therapies and technologies.

## 5. Patents

X.Y., J.X., and X.Z. are co-inventors on patents filed by N1 Life, Inc. that cover technologies discussed in the manuscript.

## Author Contributions

Conceptualization, X.Y., J.X. and X.Z.; methodology, X.Y., J.X., D.S, and X.Z.; investigation, X.Y. and J.X.; validation, X.Y. and J.X.; resources, X.Y. and J.X.; data curation, X.Y. and J.X.; writing—original draft preparation, X.Y. and J.X.; writing—review and editing, D.S. and X.Z.; visualization, X.Y. and J.X.; supervision, X.Z.; project administration, J.X.; funding acquisition, X.Z. All authors have read and agreed to the published version of the manuscript.

## Data Availability Statement

The data presented in this study are available on request from the corresponding author.

## Acknowledgments

The authors thank Suya Biotech CO., LTD. for performing and validating *in vitro* studies, and Glow Biosciences LLC for performing *in vivo* studies. Special thanks to Dr. Jieliang Wang and Dr. Jiuzhi Sun for the discussion.

