## Supplementary Material for "Efficient mRNA Delivery *In Vitro* and *In Vivo* Using a Polycharged Biodegradable Nanomaterial"

### Table of Contents

**Figure S1.** Phase contrast and GFP fluorescent microscopy images of 18 cell lines transfected with N1-501/eGFP mRNA or Lipofectamine 3000/eGFP mRNA nanocomplexes. Pictures were captured 24 hours post-transfection (Scale bar, 400 µm)..........S3

**Figure S2.** Mean fluorescence intensities (MFI) of 18 cell lines treated with N1-501/eGFP mRNA and Lipofectamine 3000/eGFP mRNA complexes at a mRNA dose of 60 ng and N1-501 dose of 0.30 µL per well. The fluorescence images were captured at 4, 12, and 24 hours post-transfection. The MFI was calculated by dividing the total fluorescence by the number of GFP-positive cells..........S6

**Figure S3.** The correlation between cell viability and transfection efficiency was analyzed across 18 cell lines transfected with (A) N1-501/eGFP mRNA and (B) Lipofectamine 3000/eGFP mRNA nanoparticles for 24 hours. The summary statistics are shown in the tables below..........S7

**Figure S4.** Cell viability of HEK 293T treated with N1-501/eGFP mRNA nanoparticles formulated in different buffers, pH range, and media. Viability was evaluated using CCK-8 assay after 24 hours of treatment. ..........S8

**Figure S5.** The shelf life of N1-501 was evaluated by transfection efficacy *in vitro*. HEK 293T cell viability was evaluated 24 hours after treatment with N1-501/eGFP mRNA nanoparticles using CCK-8 assay. ..........S9

**Table S1.** Cell culture and seeding densities for the 18 cell lines evaluated in this study..........S10

**Table S2.** Cell viability of 18 cells treated with N1-501/eGFP mRNA, and Lipofectamine 3000/eGFP mRNA complexes for 24 hours, compared to negative control groups. All studies were conducted using an mRNA dose of 60 ng per well and evaluated with a CCK-8 assay..........S11

**Table S3.** Nanoparticle size, polydispersity, and zeta potential of N1-501/eGFP mRNA measured by Malvern Zetasizer Pro..........S12

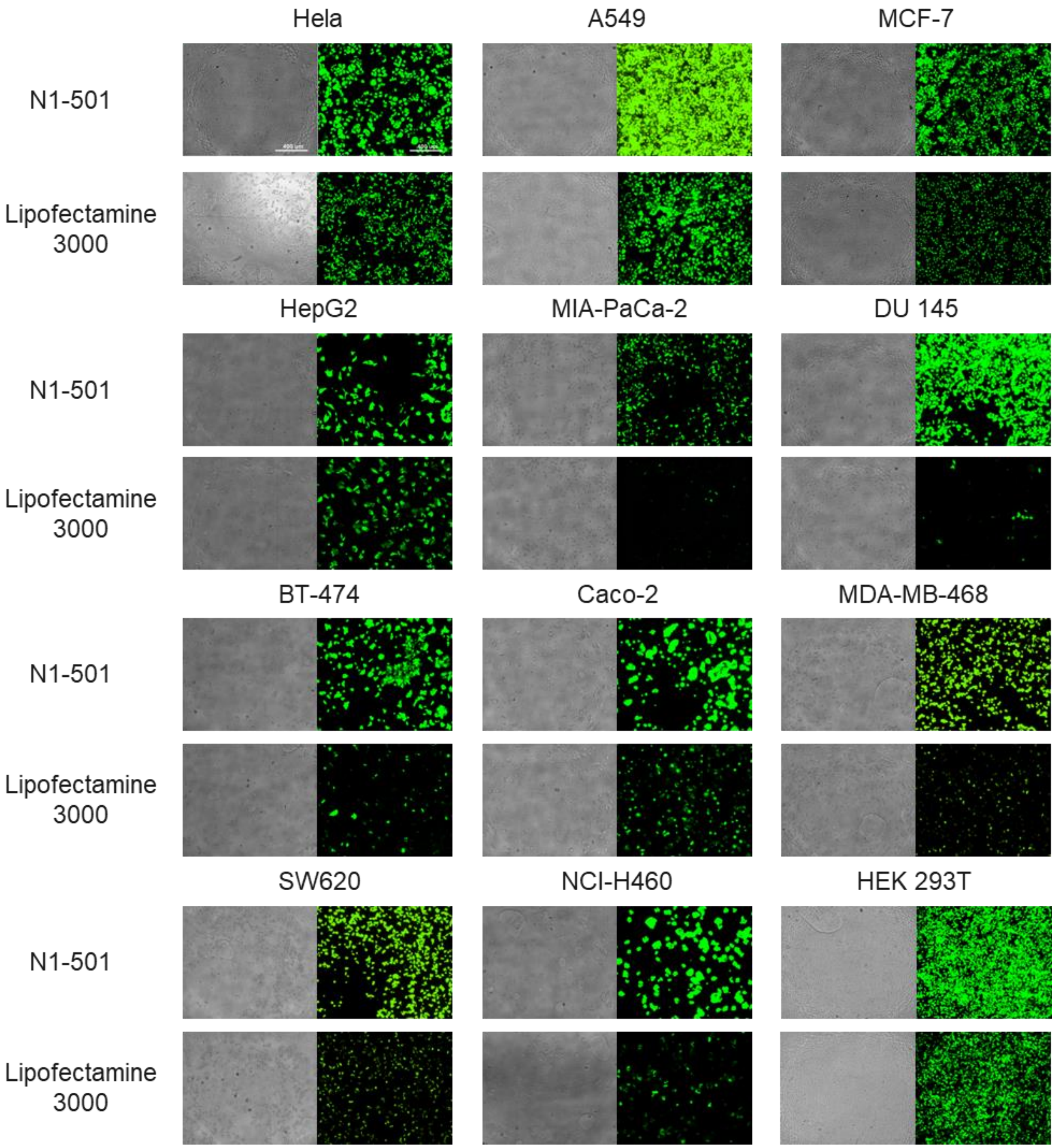
Hela
A549
MCF-7
N1-501
Lipofectamine 3000
HepG2
MIA-PaCa-2
DU 145
N1-501
Lipofectamine 3000
BT-474
Caco-2
MDA-MB-468
N1-501
Lipofectamine 3000
SW620
NCI-H460
HEK 293T
N1-501
Lipofectamine 3000

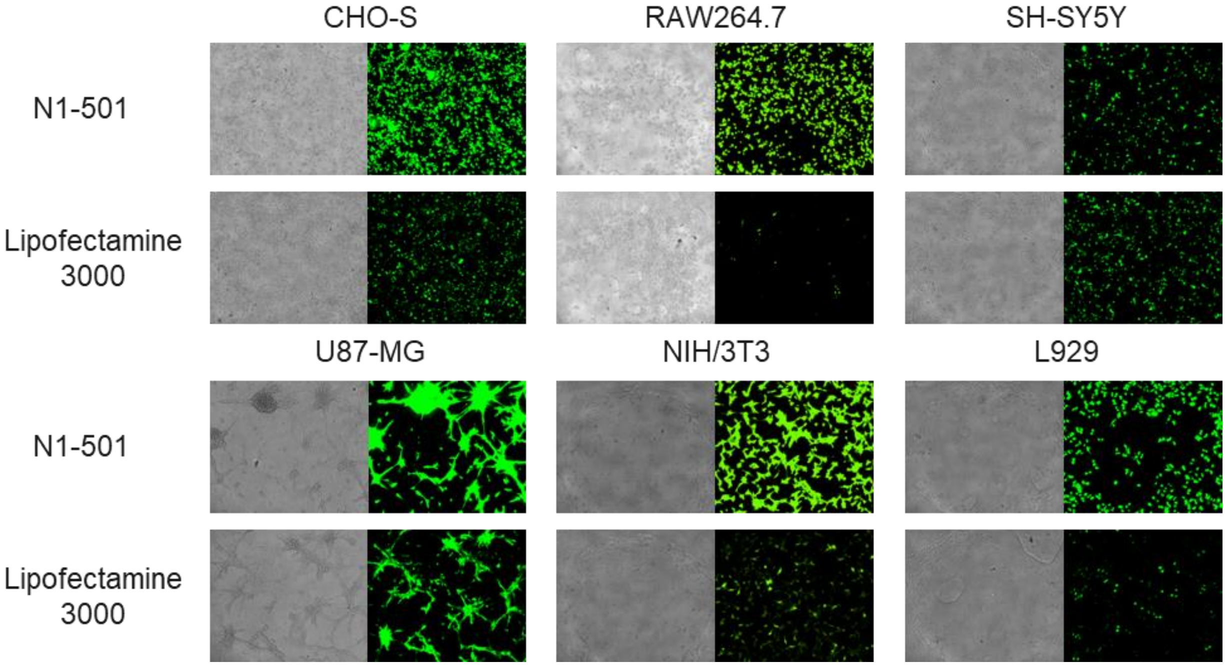


**Figure S1.** Phase contrast and GFP fluorescent microscopy images of 18 cell lines transfected with N1-501/eGFP mRNA or Lipofectamine 3000/eGFP mRNA nanocomplexes. Pictures were captured 24 hours post-transfection (Scale bar, 400 μm).

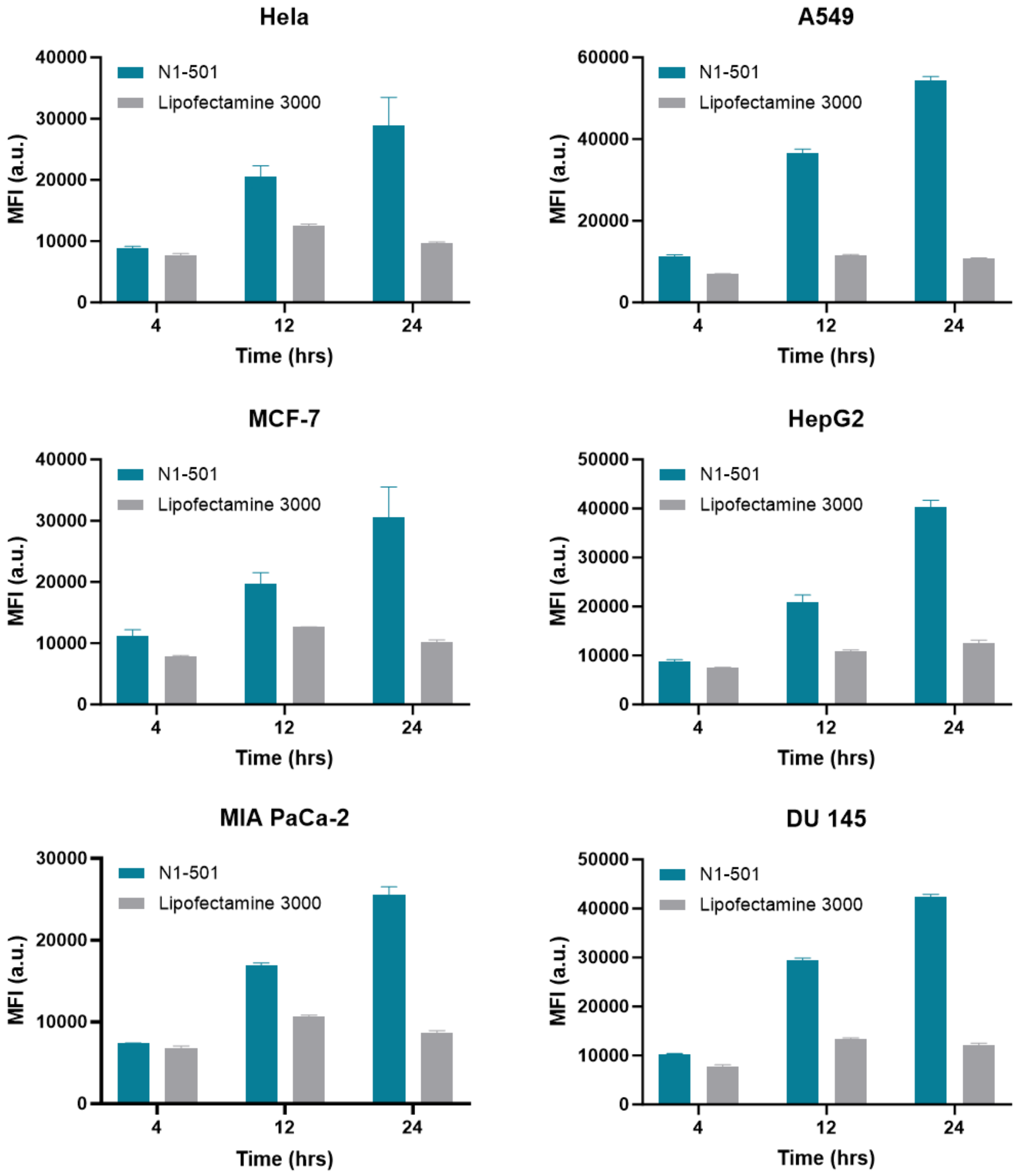
Hela
A549
MCF-7
HepG2
MIA PaCa-2
DU 145
N1-501
Lipofectamine 3000
MFI (a.u.)
Time (hrs)
4
12
24
0
10000
20000
30000
40000
50000
60000

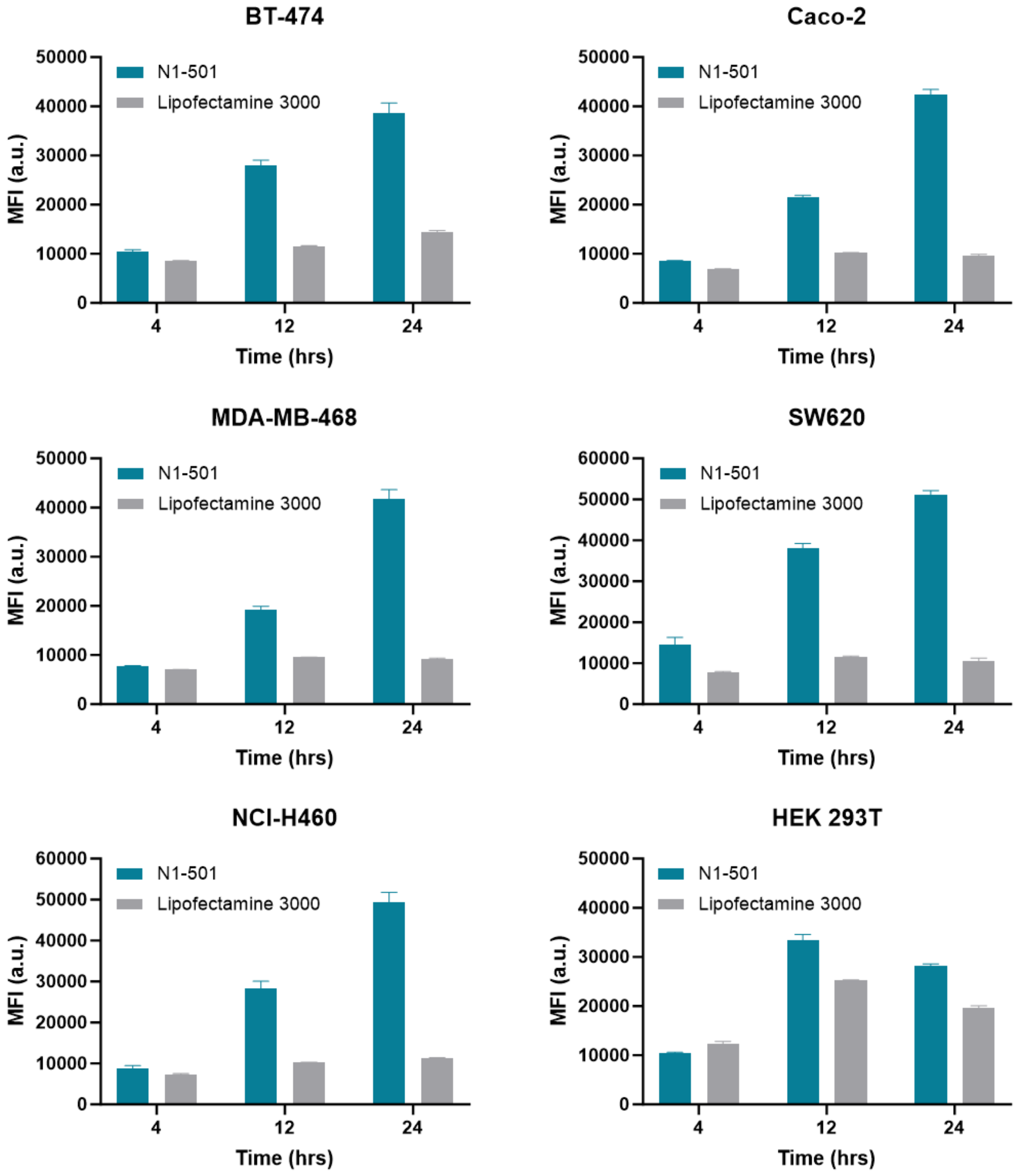
BT-474
Caco-2
MDA-MB-468
SW620
NCI-H460
HEK 293T
N1-501
Lipofectamine 3000
MFI (a.u.)
Time (hrs)
4
12
24
0
10000
20000
30000
40000
50000
60000

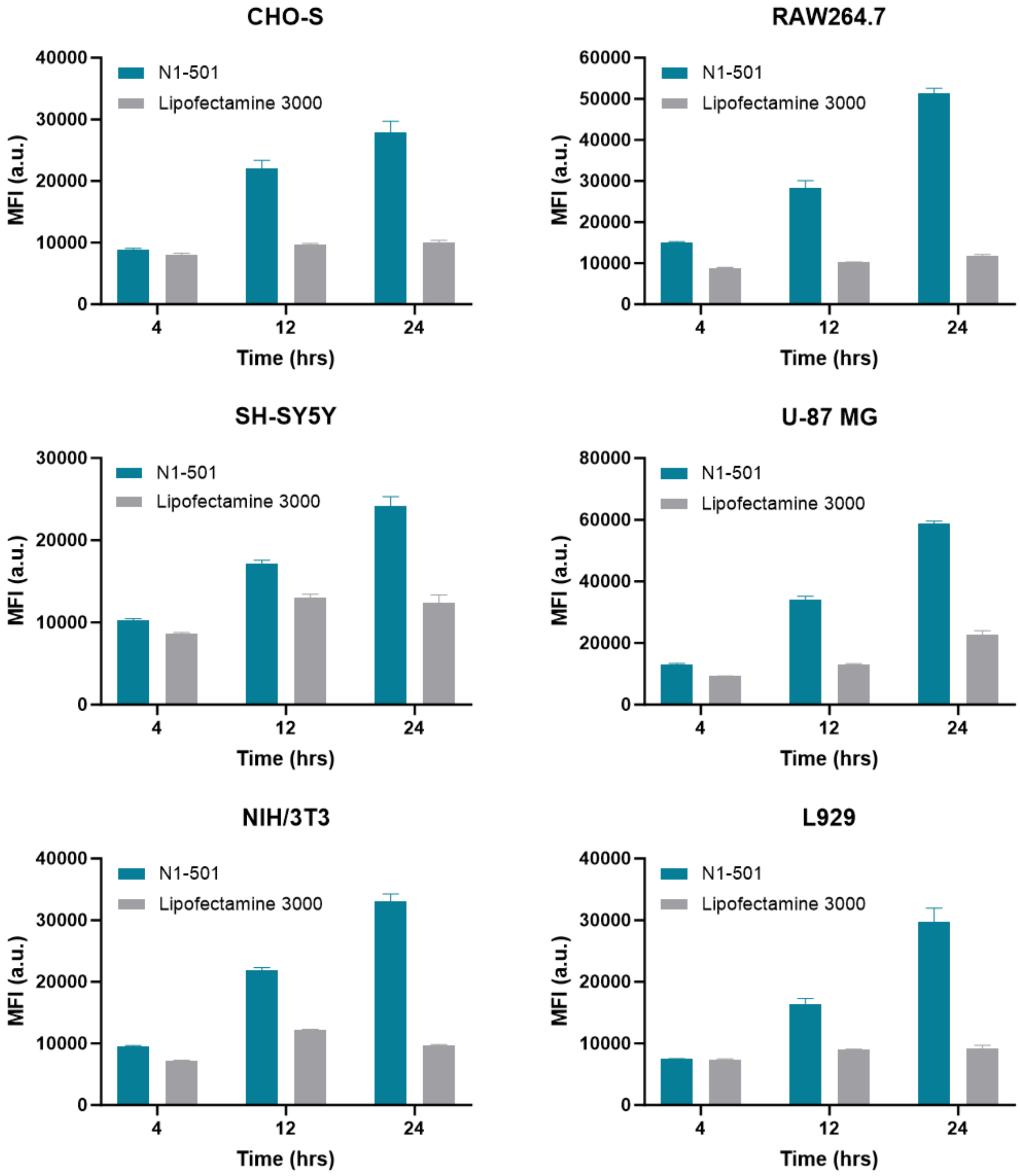


**Figure S2.** Mean fluorescence intensities (MFI) of 18 cell lines treated with N1-501/eGFP mRNA and Lipofectamine 3000/eGFP mRNA complexes at a mRNA dose of 60 ng and N1-501 dose of 0.30 µL per well. The fluorescence images were captured at 4, 12, and 24 hours post-transfection. The MFI was calculated by dividing the total fluorescence by the number of GFP-positive cells.

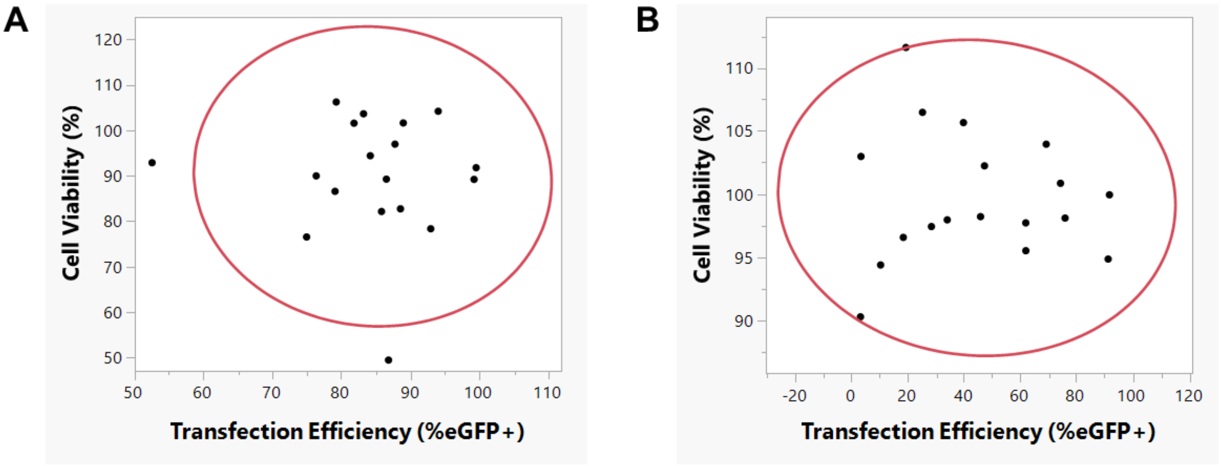


**Summary Statistics** (A)

| | Value | Lower 95% | Upper 95% | Signif. Prob |
|---|---|---|---|---|
| Correlation | -0.03673 | -0.49511 | 0.437647 | 0.8850 |
| Covariance | -5.24619 | | | |
| Count | 18 | | | |

**Summary Statistics** (B)

| | Value | Lower 95% | Upper 95% | Signif. Prob |
|---|---|---|---|---|
| Correlation | -0.04554 | -0.50175 | 0.43048 | 0.8576 |
| Covariance | -6.71042 | | | |
| Count | 18 | | | |

**Figure S3.** The correlation between cell viability and transfection efficiency was analyzed across 18 cell lines transfected with (A) N1-501/eGFP mRNA and (B) Lipofectamine 3000/eGFP mRNA nanoparticles for 24 hours. The summary statistics are shown in the tables below.

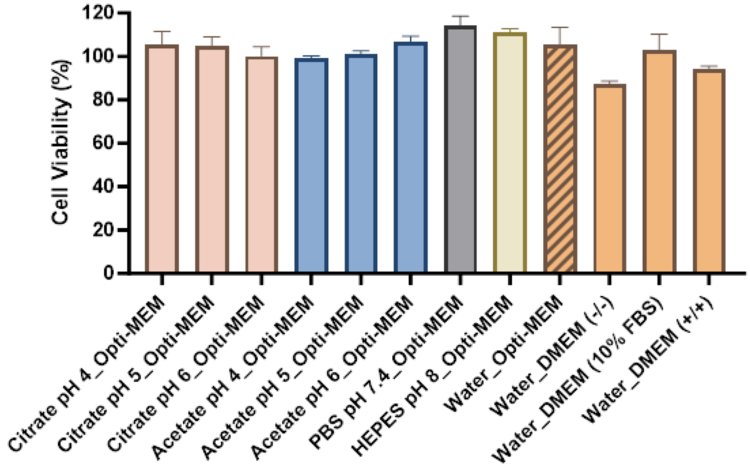


**Figure S4.** Cell viability of HEK 293T treated with N1-501/eGFP mRNA nanoparticles formulated in different buffers, pH range, and media. Viability was evaluated using CCK-8 assay after 24 hours of treatment.

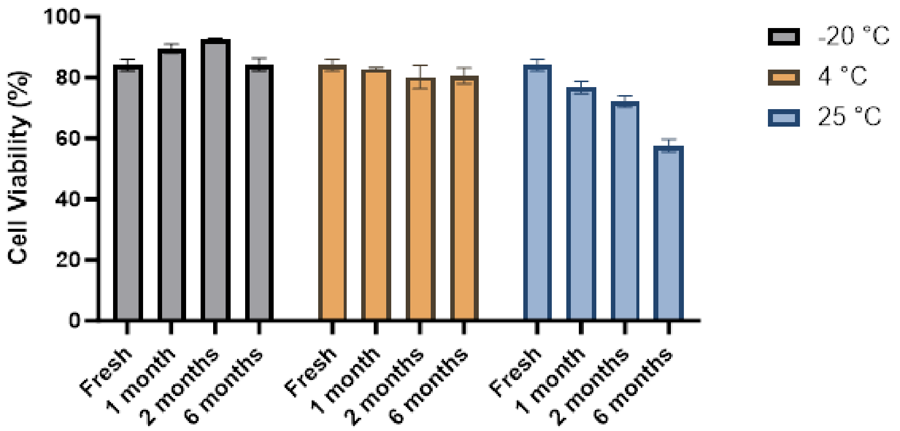


**Figure S5.** The shelf life of N1-501 was evaluated by transfection efficacy *in vitro*. HEK 293T cell viability was evaluated 24 hours after treatment with N1-501/eGFP mRNA nanoparticles using CCK-8 assay.

**Table S1.** Cell culture and seeding densities for the 18 cell lines evaluated in this study.

| Cell name | Source | Culture medium | Growth properties | Seeding density/well |
|---|---|---|---|---|
| HeLa | Nanjing RegeneCore Biotech Co.,Ltd | DMEM+10%FBS | adherent | 10,000 |
| HEK 293T | Cell Bank/Stem Cell Bank, Chinese Academy of Sciences | DMEM+10%FBS | adherent | 15,000 |
| A549 | Cell Bank/Stem Cell Bank, Chinese Academy of Sciences | DMEM+10%FBS | adherent | 7,500 |
| MCF-7 | Vazyme Biotech Co.,Ltd | DMEM+10%FBS | adherent | 10,000 |
| HepG2 | Nanjing Medical University | DMEM+10%FBS | adherent | 7,500 |
| NIH/3T3 | Nanjing University | DMEM+10%FBS | adherent | 7,500 |
| L929 | Zhongmei Guanke Biology Technology Co., Ltd | DMEM+10%FBS | adherent | 7,500 |
| MIA PaCa-2 | Cell Bank/Stem Cell Bank, Chinese Academy of Sciences | DMEM+10%HI FBS+2.5%HS | adherent | 7,500 |
| DU 145 | Cell Bank/Stem Cell Bank, Chinese Academy of Sciences | DMEM+10%FBS+1%NEAA | adherent | 7,500 |
| BT-474 | Jiangsu KeyGEN BioTECH Corp., Ltd | RPMI-1640+10%FBS | adherent | 15,000 |
| Caco-2 | FuHeng BioLogy | RPMI-1640+10%FBS | adherent | 15,000 |
| MDA-MB-468 | Cell Bank/Stem Cell Bank, Chinese Academy of Sciences | RPMI-1640+10%FBS | adherent | 17,500 |
| SW620 | Cell Bank/Stem Cell Bank, Chinese Academy of Sciences | RPMI-1640+10%FBS | adherent | 15,000 |
| NCI-H460 | Nanjing Cobioer Biosciences CO.,LTD | RPMI-1640+10%FBS | adherent | 6,000 |
| RAW264.7 | Jiangsu Kanion Pharmaceutical Co., Ltd | DMEM+10%HI FBS | adherent | 25,000 |
| SH-SY5Y | Cell Bank/Stem Cell Bank, Chinese Academy of Sciences | DMEM/F12(1:1)+10%FBS | adherent | 25,000 |
| U-87 MG | Nanjing RegeneCore Biotech Co.,Ltd | MEM+10%FBS+1%NEAA+1 mM NaP | adherent | 15,000 |
| CHO-S | Nanjing RegeneCore Biotech Co.,Ltd | RPMI-1640+10%FBS | non-adherent | 15,000 |

*Abbreviations: Dulbecco's Modified Eagle’s Medium (DMEM), Minimum Essential Media (MEM), Ham's F-12 Nutrient Mixture (F12), RPMI-1640 Medium (RPMI-1640), bovine serum (FBS), heat-inactivated FBS (HI FBS), horse serum (HS), non-essential amino acids (NEAA), sodium pyruvate (NaP).

**Table S2.** Cell viability of 18 cells treated with N1-501/eGFP mRNA, and Lipofectamine 3000/eGFP mRNA complexes for 24 hours, compared to negative control groups. All studies were conducted using an mRNA dose of 60 ng per well and evaluated with a CCK-8 assay.

| Cell name | Cell Viability (%) | | | |
|---|---|---|---|---|
| | Untreated cell | eGFP mRNA only | Lipofectamine 3000 | N1-501 |
| Hela | 100.00±2.18 | 100.64±4.48 | 97.73±2.93 | 103.76±3.82 |
| A549 | 100.00±14.14 | 98.25±10.81 | 99.95±14.68 | 91.90±11.67 |
| MCF-7 | 100.00±1.03 | 103.13±2.07 | 102.24±1.23 | 101.70±1.33 |
| HepG2 | 100.00±5.97 | 99.38±6.65 | 98.11±6.99 | 97.08±6.86 |
| MIA PaCa-2 | 100.00±8.34 | 108.92±8.54 | 111.60±10.43 | 89.37±6.57 |
| DU 145 | 100.00±8.03 | 103.78±9.85 | 102.98±10.23 | 101.75±11.17 |
| BT-474 | 100.00±16.25 | 90.30±12.83 | 94.40±14.44 | 90.09±17.55 |
| Caco-2 | 100.00±5.09 | 102.17±5.11 | 98.22±6.51 | 76.64±4.24 |
| MDA-MB-468 | 100.00±6.03 | 98.37±7.12 | 97.97±6.57 | 86.69±4.25 |
| SW620 | 100.00±4.48 | 97.83±3.07 | 95.53±4.37 | 82.84±4.06 |
| NCI-H460 | 100.00±6.88 | 98.44±6.71 | 97.44±4.48 | 82.25±5.79 |
| HEK 293T | 100.00±3.16 | 100.26±2.71 | 94.87±2.38 | 89.32±2.78 |
| CHO-S | 100.00±2.93 | 109.40±4.20 | 105.66±4.35 | 104.32±5.98 |
| RAW264.7 | 100.00±8.66 | 100.14±9.69 | 90.30±7.72 | 49.54±6.69 |
| SH-SY5Y | 100.00±8.23 | 98.50±20.96 | 103.95±10.48 | 93.00±15.18 |
| U-87 MG | 100.00±23.63 | 96.48±19.71 | 100.86±22.82 | 106.36±8.65 |
| NIH/3T3 | 100.00±3.88 | 101.65±6.17 | 106.47±5.78 | 94.53±6.98 |
| L929 | 100.00±8.14 | 94.38±6.86 | 96.58±6.68 | 78.45±8.07 |

**Table S3.** Nanoparticle size, polydispersity, and zeta potential of N1-501/eGFP mRNA measured by Malvern Zetasizer Pro.

| Sample name | N1-501/eGFP mRNA |
|---|---|
| Size (nm) | 119.6±13.1 |
| PDI | 0.20±0.08 |
| Zeta Potential (mV) | 9.1±3.2 |
